# Pharmacological and genetic manipulations of Ca^2+^ signaling have contrasting effects on auxin-regulated trafficking

**DOI:** 10.1101/2020.09.04.283101

**Authors:** Ren Wang, Ellie Himschoot, Matteo Grenzi, Jian Chen, Melanie Krebs, Karin Schumacher, Moritz K. Nowack, Daniël Van Damme, Ive De Smet, Tom Beeckman, Alex Costa, Jiří Friml, Steffen Vanneste

**Affiliations:** Ghent University, Department of Plant Biotechnology and Bioinformatics, 9052 Ghent, Belgium; VIB Center for Plant Systems Biology, 9052 Ghent, Belgium; Department of Biosciences, University of Milan, 20133 Milan, Italy; Centre for Organismal Studies, Plant Developmental Biology, University of Heidelberg, 69120 Heidelberg, Germany; Institute of Biophysics, National Research Council of Italy (CNR), 20133 Milano, Italy; Institute of Science and Technology Austria (IST Austria), 3400 Klosterneuburg, Austria; Lab of Plant Growth Analysis, Ghent University Global Campus, Incheon 21985, Republic of Korea

**Keywords:** Auxin, Calcium, Trafficking, Signaling

## Abstract

A large part of a plants’ developmental plasticity relies on the activities of the phytohormone auxin and the regulation of its own distribution. This process involves a cohort of transcriptional and non-transcriptional effects of auxin on polar auxin transport, regulating the abundancy, biochemical activity and polar localization of the molecular components, predominantly PIN auxin exporters. While the transcriptional auxin signaling cascade has been well characterized, the mechanism and role of non-transcriptional auxin signaling remains largely elusive. Here, we addressed the potential involvement of auxin-induced Ca^2+^ signaling in auxin’s inhibitory effect on PIN endocytic trafficking. On the one hand, exogenous manipulations of Ca^2+^ availability and signaling effectively antagonized auxin effects suggesting that auxin-induced Ca^2+^ signaling is required for inhibition of internalization. On the other hand, we addressed the auxin-mediated inhibition of PIN internalization in the auxin signaling (*tir1afb2,3*) or Ca^2+^ channel (*cngc14*) mutants. These mutants were strongly defective in auxin-triggered Ca^2+^ signaling, but not in auxin-inhibited internalization. These data imply that, while Ca^2+^ signaling may be required for normal PIN trafficking, auxin-mediated increase in Ca^2+^ signaling is not a direct part of a downstream mechanism that mediates auxin effects on Brefeldin A-visualized PIN intercellular aggregation. These contrasting results obtained by comparing the mutant analysis versus the exogenous manipulations of Ca^2+^ availability and signaling illustrate the critical importance of genetics to unravel the role of Ca^2+^ in a process of interest.

## INTRODUCTION

Calcium cross-links pectinate polymers and is therefore an important structural determinant of the cell wall (Feng et al., 2018; Cosgrove and Anderson, 2020), and inside the cell it directly modulates the biochemical activities of proteins, Ca^2+^ sensors (relays and responders) and phospholipids, impacting on numerous cellular processes (Himschoot et al., 2017; Kudla et al., 2018). The pleiotropic activities of Ca^2+^ necessitate submicromolar range Ca^2+^ levels in the cytoplasm, while in the apoplast and in the lumen of organelles, Ca^2+^ levels are several orders of magnitude higher (Stael et al., 2012; Costa et al., 2018). Such steep concentration gradients allow to trigger a local significant increase in Ca^2+^ levels by the simple opening of a few channels in response to a specific stimulus (Demidchik et al., 2018). In a typical signaling cascade, the cytoplasmic increase of Ca^2+^ is decoded by specialized proteins that translate the Ca^2+^ signal into defined cellular responses, such as the modulation of channels or kinases (Kudla et al., 2018).

The plant Ca^2+^ signaling toolkit is strongly diversified in comparison to the one in animals (Edel et al., 2017), most prominently reflected in the existence of plant and animal specific Ca^2+^ signaling components, such as inositol (1,4,5)-triphosphate receptors, and ryanodine receptors. Despite this important diversification, the current commonly used Ca^2+^ pharmacology consists of very general inhibitors or chelators, or inhibitors that were designed to target mammalian Ca^2+^ channels and signaling components (De Vriese et al., 2018). In most cases, the molecular targets of the inhibitors are not well enough conserved or even absent in plants, making it difficult to make strong claims based on inhibitor studies.

The signaling function of Ca^2+^ is currently best understood in the context of abiotic and biotic stress responses (Kudla et al., 2018), guard cell movement (Konrad et al., 2018) and in pollen tubes (Guo and Yang, 2020). In contrast, the function of the since long described auxin-induced Ca^2+^ response remains largely elusive (Vanneste and Friml, 2013; Shih et al., 2015; Dindas et al., 2018). Only recently, this for long overlooked aspect of auxin signaling regained attention with the identification of the non-selective cation channel CNGC14 as a critical component of auxin-induced Ca^2+^ entry (Shih et al., 2015; Dindas et al., 2018). CNGC14 activity was proposed to participate in root gravitropism (Shih et al., 2015) and root hair development (Dindas et al., 2018; Brost et al., 2019). Additionally, manipulations of Ca^2+^ availability and channels revealed connections to polar auxin transport (Dela Fuente and Leopold, 1973) and polarization of auxin transporters (Zhang et al., 2011; Li et al., 2019), indicating an important interplay between auxin and Ca^2+^.

In contrast to our poor understanding of auxin-induced Ca^2+^ signaling, the mechanism of auxin-induced transcriptional changes has been characterized in great detail (Lavy and Estelle, 2016; Roosjen et al., 2018; Powers and Strader, 2020). The canonical pathway for auxin-induced transcription involves the auxin-stabilized interaction between TIR1/AFB F-box proteins and Aux/IAA transcriptional corepressors (Dharmasiri et al., 2005; Kepinski and Leyser, 2005). This results in the ubiquitination and proteolysis of the latter (Nemhauser, 2018). Consequently, the transcriptional repressive effect imposed by Aux/IAA on ARF transcription factors is released, and auxin-induced transcription can proceed (Pierre-Jerome et al., 2016; Roosjen et al., 2018). Recently, a non-transcriptional branch of TIR1-based auxin signaling was demonstrated to effect acute inhibition of elongation (Fendrych et al., 2018; Gallei et al., 2020). Moreover, the non-transcriptional repertoire of TIR1/AFBs was recently further expanded by the observation that *tir1/afb2,3* mutants are defective in auxin-induced Ca^2+^ signaling (Dindas et al., 2018). Additionally, auxin signals converging on pavement cell morphogenesis (Xu et al., 2010), lipid composition and distribution (Pan et al., 2009; Li et al., 2015; Platre et al., 2019), cell division during lateral root formation (Huang et al., 2019), suppression of auxin biosynthesis (Wang et al., 2020), stability and polarity of PIN proteins (Abas et al., 2006; Sauer et al., 2006; Baster et al., 2013; Prat et al., 2018; Mazur et al., 2020), and their internalization (Paciorek et al., 2005; Robert et al., 2010; Platre et al., 2019), possibly act via alternative auxin perception mechanisms, such as the receptor like kinase family TRANSMEMBRANE KINASE1-4 (Cao et al., 2019; Huang et al., 2019; Platre et al., 2019).

Based on the non-transcriptional character of the inhibition of internalization by NAA (Robert et al., 2010; Zhang et al., 2020), we postulated that auxin-induced Ca^2+^ signaling could be a signaling component in this response to NAA. Therefore, followed two strategies. On the one hand, we manipulated NAA-induced Ca^2+^ signaling using inhibitors or washing seedlings in Ca^2+^ free medium. We validated their effects on NAA-induced Ca^2+^ signaling, and assessed their impact on NAA-inhibited internalization. This approach provided a convincing and tight correlation between NAA-induced Ca^2+^ signaling and NAA’s ability to inhibit internalization. However, this correlation between Ca^2+^ signaling and inhibition of internalization could not be confirmed in *tir1/afb* and *cngc14*, two mutants that are specifically defective auxin-induced Ca^2+^ signaling. This discrepancy in outcome between both approaches calls for extreme caution when analyzing the role of Ca^2+^ in a process of interest as current pharmacology or manipulations of Ca^2+^ availability are prone to pleiotropic, misleading effects.

## RESULTS

### Characterization of NAA-induced Ca^2+^ signaling

The synthetic auxin, 1-naphthaleneacetic acid (NAA) is widely used in auxin biology as a proxy for the endogenous auxin indole-3-acetic acid (IAA) and is a more potent inhibitor of internalization than IAA (Paciorek et al., 2005). Because we wanted to evaluate the role of Ca^2+^ signaling in inhibition of internalization, we set out to characterize the NAA-induced Ca^2+^ response in Arabidopsis root meristems in more detail. The cytoplasmic, intensiometric Ca^2+^ indicator R-GECO1 (Keinath et al., 2015) reported an instant cytosolic Ca^2+^ elevation in response to 10µM NAA application (Figure 1A,B; Supplemental Movie S1). Also at lower concentrations (1µM and 0.1µM), NAA triggered rapid Ca^2+^ signaling, albeit with smaller amplitude (Figure 1B), illustrating a dose-dependence of the maximal response, similar to the one reported for the natural auxin indole-3-acetic acid (IAA) (Dindas et al., 2018). The onset of the Ca^2+^ increase started within seconds after NAA application, and reached a maximum after ±70 sec, followed by a gradual attenuation response. Similarly, subcellular targeting of ratiometric Ca^2+^ sensors revealed a rise in Ca^2+^ concentrations in the cytoplasm (visualized with NES-YC3.6 (Krebs et al., 2012)), in the cytosol near the plasma membrane (visualized with PM-YC3.6-Lti6b (Krebs et al., 2012)), in the endoplasmic reticulum lumen (visualized with CRT-D4ER (Bonza et al., 2013)) and in mitochondria (visualized with 4mt-YC3.6 (Loro et al., 2012)) (Supplemental Figure S1A-D). The signals in the ER and mitochondria were slightly delayed compared to the other reporters (Supplemental Figure S1C,D), suggesting that these organelles may act as Ca^2+^ sinks for attenuation of the cytoplasmic Ca^2+^ signal. The lack of Ca^2+^ response after benzoic acid (BA) treatment (Supplemental Figure S1E,G), shows that the Ca^2+^ response to NAA was not a response to acidification associated with NAA treatment. The NAA-induced Ca^2+^ response could be inhibited pharmacologically by the Ca^2+^ channel inhibitors Bepridil and Nifedipine (De Vriese et al., 2018; De Vriese et al., 2019) (Supplemental Figure S1F,G; Supplementary Movies S2,S3).

**Figure 1.**
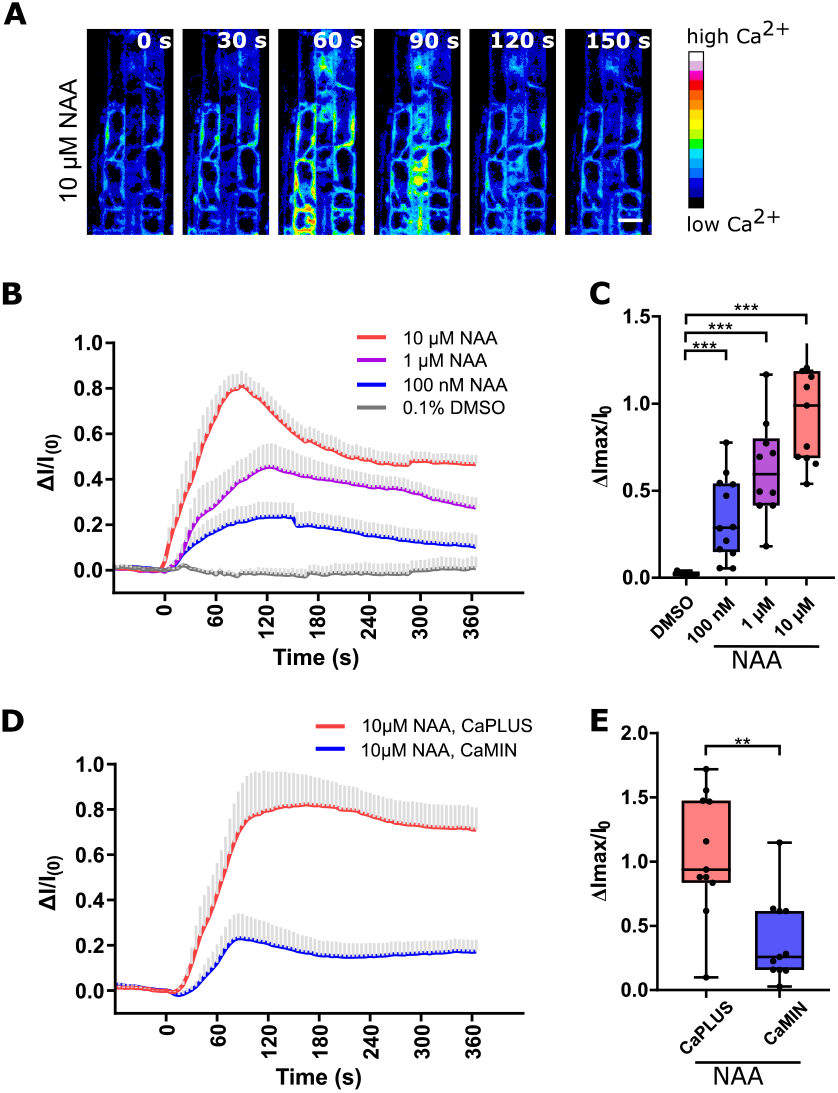
Apoplastic Ca^2+^ determines the amplitude of auxin-induced cytosolic Ca^2+^ dynamics. **A**. Single frames of the dynamic response of the Ca^2+^ sensor, R-GECO1, at indicated time points after 10µM NAA treatment. Scale bar = 20 µm. **B**. Averaged and normalized R-GECO1 fluorescence intensities over time upon treatment with 0.1% DMSO, 10μM NAA, 1μM NAA, or 100nM NAA. (nr of seedlings = 3, 12, 10, 12 respectively, means +s.e.m.). DMSO and NAA treatments were applied at time point 0. **C**. Boxplot representation of the maximal amplitude of the treatments described in **B**. **D**. The averaged normalized R-GECO1 fluorescence intensities over time upon treatment with 10μM NAA following a 30min pretreatment with CaPLUS or CaMIN (nr of seedlings = 11; 3 repeats; means +s.e.m.). NAA treatments were applied at time point 0s. **E**. Boxplot representation of the maximal amplitude of the treatments described in **D**. For all box plots, the central line indicates the median, the bottom and top edges of the box indicate the interquartile range. The box plot whiskers are plotted down to the minimum and up to the maximum value. Data were analyzed by an unpaired two-tailed t-test with Welch correction. **P < 0.01, ***P < 0.001.

Additionally, we modulated the available extracellular Ca^2+^ by washing the seedlings with 0.5xMS medium that lacked CaCl_2_ (hereafter referred to as CaMIN). This simple treatment was sufficient to reduce the NAA-induced Ca^2+^ response in comparison to normal 0.5xMS medium (≈1.5mM CaCl_2_, hereafter CaPLUS) (Figure 1C). Jointly, these data suggest that NAA triggers a complex Ca^2+^ response that largely depends on extracellular Ca^2+^.

### NAA-mediated Inhibition of internalization correlates with Ca^2+^ signaling

The synthetic auxin NAA interferes rapidly via a non-transcriptional pathway with the internal accumulation of plasma membrane proteins (hereafter referred to as internalization) in Brefeldin A (BFA)-induced intracellular endosomal aggregates (so-called BFA bodies) (Paciorek et al., 2005; Robert et al., 2010). This effect of NAA was concentration dependent, showing the maximum inhibitory effect at 10µM (Supplemental Figure S2A,B), correlating with the dose-dependence of the maximal NAA-induced Ca^2+^ response (Figure 1C,D). Given the immediacy of both processes, we postulated that auxin-induced Ca^2+^ responses reflects an auxin signaling cascade involved in NAA-regulated internalization.

Indeed, when we used the organic Ca^2+^ channel blockers Nifedipine and Bepridil at concentrations that interfered with NAA-induced Ca^2+^ entry (Supplemental Figure 1F,G), PIN1 internalization was restored in BFA/NAA co-treated roots (Supplemental Figure S2C). A similar nullification on NAA-inhibited PIN1 internalization was achieved using the membrane-permeable calmodulin inhibitor W-7 (Supplemental Figure S2C). This suggests that Ca^2+^ increase and its downstream signaling is required for NAA’s inhibitory effect on PIN1 internalization. Given the potential off-target effects of the Ca^2+^ channel inhibitors (De Vriese et al., 2018), we also evaluated the effect of CaMIN on NAA-inhibited PIN1 internalization. Similarly to Bepridil, Nifedipine and W-7, a 30min CaMIN pretreatment was sufficient to restore PIN internalization in BFA/NAA co-treated roots (Figure 2A,B). To evaluate the specificity of the treatment to Ca^2+^ availability, we analyzed PIN1 internalization in CaMIN supplemented with either 1.5mM CaCl_2_ (comparable to 0.5xMS) or 1.5mM MgCl_2_ (Figure 2A,B). The addition of CaCl_2_ fully restored the NAA sensitivity of PIN1 internalization. In contrast, the internalization in roots treated with MgCl_2_ supplemented CaMIN could not restore the NAA sensitivity, indicating the specificity of Ca^2+^ in this process. This effect of CaMIN on NAA-inhibited internalization could also be observed for other plasma membrane cargoes such as PIN2, AUX1 in the protophloem and NPSN12 (WAVE131-YFP) (Figure 2C-F). Jointly, these findings strongly support a notion that Ca^2+^ is required for NAA’s inhibitory effect on internalization of plasma membrane proteins.

**Figure 2.**
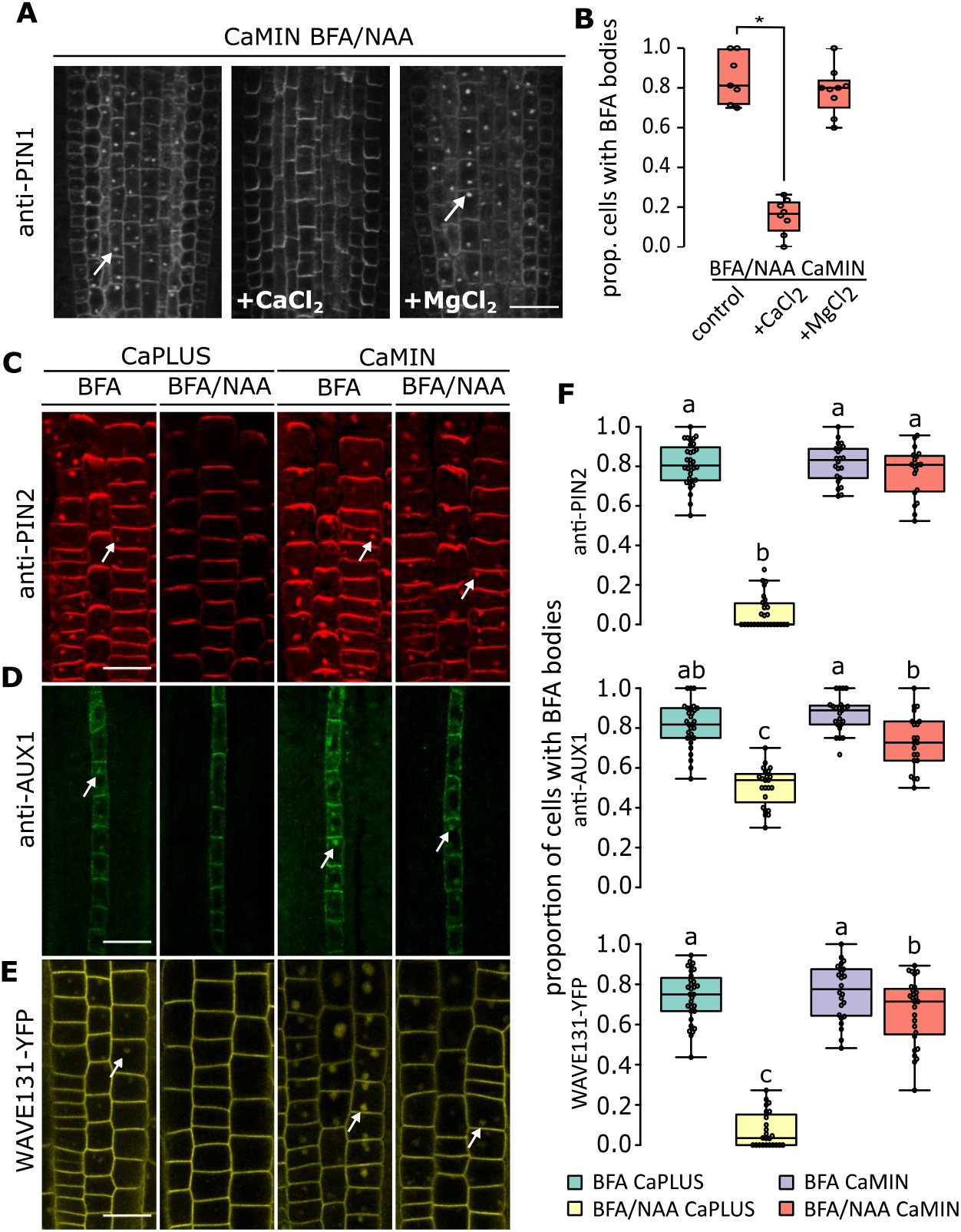
NAA inhibition of internalization requires extracellular Ca^2+^. **A**. Immunolocalisation of PIN1 in 3 day-old seedling roots pretreated with CaMIN (30min), followed by BFA and NAA (co)treatment (1h), in CaMIN, and in CaMIN supplemented with either 1.5 mM CaCl_2_ and CaMIN supplemented with 1.5mM MgCl_2_. Scale bar = 20μm. **B**. Quantification of the proportion of cells that have PIN1 in BFA bodies for the conditions in **A**, and corresponding controls in CaPLUS.(n ≥ 3 per treatment per repeat; 2 independent repeats). For all box plots, the central line indicates the median, the bottom and top edges of the box the interquartile range, and the box plot whiskers are plotted down to the minimum and up to the maximum value. Data were analyzed using a logistic regression model. * indicates P ≤ 0.05, Wald-type test. **C**. Immunolocalization of PIN2 in 3 day-old seedling roots after BFA or BFA/NAA treatments in CaPLUS and CaMIN conditions. **D**. Immunolocalization of AUX1 in 3 day-old seedling roots after BFA or BFA/NAA treatments in CaPLUS and CaMIN conditions. **E**. WAVE131(NPSN12)-YFP localisation in 3 day-old seedling roots after BFA or BFA/NAA treatments in CaPLUS and CaMIN conditions. Concentrations used for PIN2 and WAVE131-YFP BFA: 25μM, NAA: 10μM, 1h; Concentrations used for AUX1 BFA: 50μM, NAA: 10μM, 90min. **F**. Quantification of the proportion of cells that have PIN2 (n=32;28;22;19 roots in total), AUX1 (n=30;26;26;22 roots in total) or WAVE131-YFP (n=31;24;24;27 roots in total) in BFA bodies, corresponding to experiments in Figure 2C-E. Center lines show the medians; box limits indicate the 25th and 75th percentiles; whiskers extend 1.5 times the interquartile range from the 25th and 75th percentiles, outliers are represented by dots. Different lowercase letters indicate significant differences (P ≤ 0.05, Wald-type test). White arrows in figures indicate proteins accumulated in BFA bodies. Scale bars represents 20 µm.

### The TIR1/AFB-CNGC14 module is not required for NAA’s inhibitory effect on internalization

TIR1/AFB-based auxin perception was demonstrated to be required for IAA-induced Ca^2+^ signaling via the Ca^2+^ permeable cation channel CNGC14 (Shih et al., 2015; Dindas et al., 2018). Assuming an auxintrigerred Ca^2+^ signal being part of the mechanism for regulation of PIN internalization, we predicted that the defective auxin-induced Ca^2+^ signaling in *tir1/afb* or *cngc14* mutants would result in NAA-insensitive PIN internalisation. Contrary to expectations based on CaMIN and Ca^2+^ signaling inhibitors, none of the tested mutants showed NAA-insensitive internalisation in CaPLUS, suggesting that the TIR1/AFB-CNGC14 Ca^2+^ signaling module is not essential for this effect. Surprisingly, PIN1 internalization was inhibited by NAA in *tir1afb1afb3, tir1afb2afb3* and three different *cngc14* alleles also in CaMIN conditions, unlike in WT and other *tir1/afb* mutants (Figure 3B-E; Supplemental Figure S3A-E). In contrast to *tir1/afb* and *cngc14* mutants, the NAA-sensitive internalization in CaMIN was not observed in the auxin biosynthesis defective mutant *yuc3,5,7,8,9* (*yucQ*) (Chen et al., 2014) (Supplemental Figure S3F). Thus, the restored NAA sensitivity in CaMIN-treated *tir1/afb* and *cncg14* mutants does not seem to be related to changes in auxin levels but is rather specific to TIR1/AFB-CNGC14-mediated auxin signaling. Jointly, these data demonstrate that in CaPLUS, NAA inhibits internalisation independently of TIR1/AFB-CNGC14-mediated auxin signaling.

**Figure 3.**
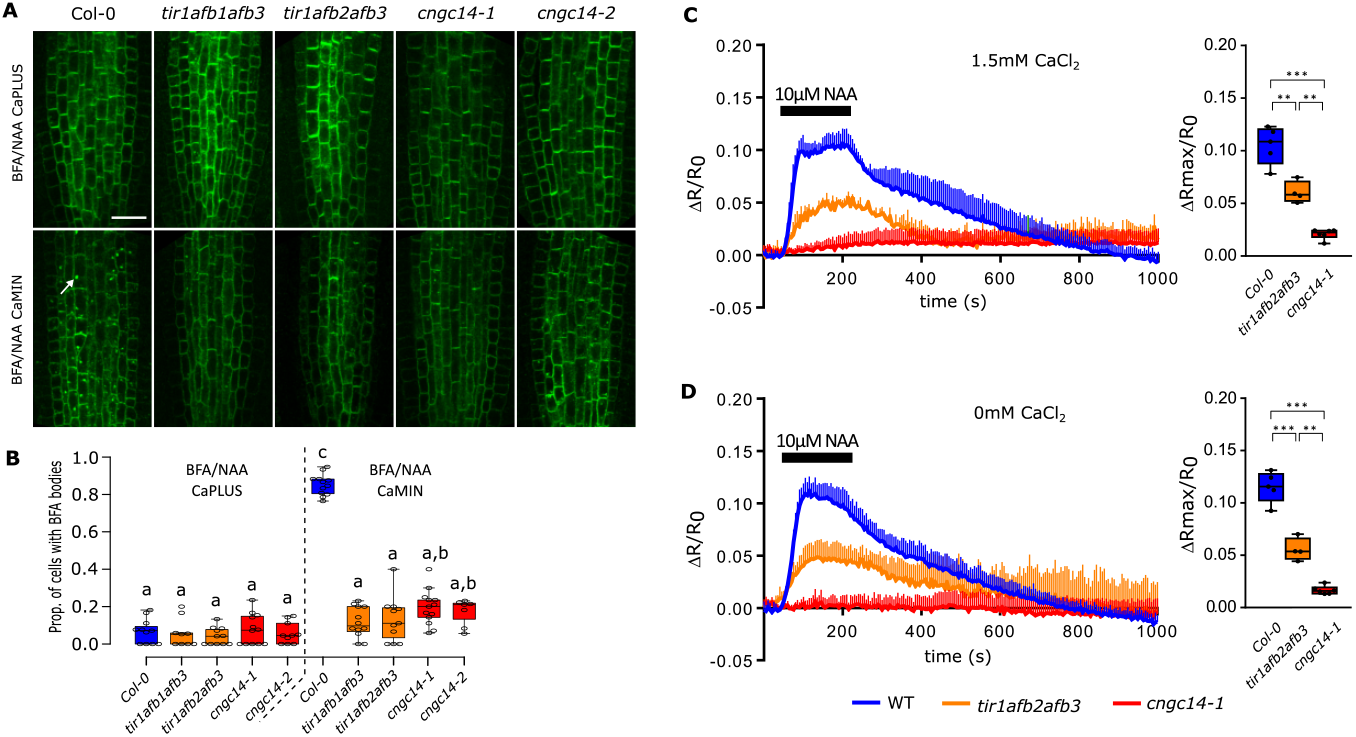
Analysis of inhibition of internalization and auxin-induced Ca^2+^ signaling in *tir1/afb* and *cngc14* mutants. **A**. Immunolocalization of PIN1 in 3 day-old roots co-treated for BFA and NAA in wild type, *tir1/afb1,3, tir1/afb2,3, cngc14-1* and *cngc14-2*, in CaPLUS and CaMIN conditions. White arrows indicate PIN1-accumulating BFA bodies. Scale bar is 20 µm. **B**. Quantification of the proportion of cells that accumulate PIN1 in BFA bodies visualized by immunolocalisation in 3 day-old roots of indicated genotypes co-treated for BFA and NAA, in CaPLUS, CaMIN as depicted in **c**. Total numbers of roots analyzed in wild type (n=11; n=12 in total), *tir1afb1afb3* (n=9; n=12 in total), *tir1afb2afb3* (n=11; n=11 in total), *cngc14-1* (n=12; n=13 in total) and *cngc14-2* (n=10; n=7 in total), representing the sum of two independent experiments. Significant differences (*P* ≤ 0.05, Wald-type test) are indicated by different lowercase letters. Center lines show the medians; box limits indicate the 25th and 75th percentiles; whiskers extend 1.5 times the interquartile range from the 25th and 75th percentiles, outliers are represented by dots. **C,D**. The Ca^2+^ dynamics (NES-YC3.60 Cameleon) in WT, *tir1afb2afb3, cngc14-1* in response to a pulse of 10µM NAA in medium containing (**C**) 1.5mM CaCl_2_ or (**D**) 0mM CaCl_2_.

The discrepancy between the exogenous Ca^2+^ manipulations and the mutant analyses, raised the hypothesis that NAA may induce Ca^2+^ signaling via a TIR1/AFB-CNGC14-independent mechanism. Therefore, we transformed the ratiometric Ca^2+^ indicator NES-YC3.6 in *tir1/afb2,3* and *cngc14-1* and analysed the Ca^2+^ responses to a pulse of NAA in these mutants. We found that the NAA-induced Ca^2+^ response was strongly reduced in *tir1afb2,3* triple mutants and was completely absent from *cngc14-1* (Figure 3A), suggesting that NAA elicits Ca^2+^ signaling through the same mechanism as IAA. Also in the absence of exogenously supplied Ca^2+^, the NAA-induced Ca^2+^ responses in *tir1afb2,3* and *cngc14-1* mutants were strongly defective (Figure 3B), excluding compensatory Ca^2+^ signaling mechanisms under low Ca^2+^ availability, as reported for pathogen-triggered immunity-associated Ca^2+^ signaling in *cngc2* and *cngc4* mutants (Tian et al., 2019). These data show that NAA-induced Ca^2+^ signaling is fully dependent on the TIR1/AFB-CNGC14 module. Notably, the YC3.6 Ca^2+^ indicator did not allow detecting the obvious differences in the NAA-induced Ca^2+^ response between CaMIN and CaPLUS that we observed when using R-GECO1 (Figure 1D). This probably reflects fundamental differences in Ca^2+^ binding properties and dynamic ranges that exist between both Ca^2+^ indicators’ properties (Nagai et al., 2004; Zhao et al., 2011; Keinath et al., 2015; Waadt et al., 2017). This also suggests that the reduction in Ca^2+^ response caused by CaMIN treatment is weaker than the one in *tir1afb2afb3* and *cngc14-1*. The normal NAA sensitivity of PIN internalisation in these mutants, that are strongly defective in auxin-induced Ca^2+^ signaling, thus provides further evidence that auxin-induced Ca^2+^ signaling is not an essential part of the mechanism by which auxin affects trafficking. This is in direct contrast to the conclusions drawn using exogenous Ca^2+^ signaling manipulations, highlighting an important lack of specificity of these treatments.

## DISCUSSION

Auxin-mediated regulation of trafficking is considered an important aspect of auxin’s self-regulating properties in plant growth and development. However, the underlying molecular mechanisms remain largely unclear. The synthetic auxins such as NAA displays a strong effect on trafficking, as illustrated by strong interference with accumulation of plasma membrane cargoes in BFA bodies (Paciorek et al., 2005). This effect was reported to be very fast, not requiring transcriptional changes and independent of canonical auxin signaling (Robert et al., 2010). Instead, an extracellular auxin perception mechanism was proposed based on the activities of AUXIN BINDING PROTEIN1 (Robert et al., 2010). Using updated genetic tools, however, the latter was put into question (Gao et al., 2015). Given the immediacy of the response, we hypothesized that auxin-induced Ca^2+^ signaling could be involved in auxin’s inhibitory effect on internalization. We established that NAA activates Ca^2+^ responses at the plasma membrane via a TIR1/AFB-CNGC14-dependent mechanism, similarly as was recently described for the endogenous auxin IAA (Shih et al., 2015; Dindas et al., 2018). Using mutants and pharmacology we interfered with auxin-induced Ca^2+^ signaling and evaluated of the impact on internalization. The pharmacological interference revealed a good correlation between intensity of auxin-induced Ca^2+^ signaling and NAA’s ability to inhibit internalization, supporting our original notion. In striking contrast, genetic disruption of auxin-induced Ca^2+^ signaling did not affect NAA’s ability to inhibit internalization. These data unequivocally show that auxin-induced Ca^2+^ signaling does not inhibit internalization. It does however, not exclude roles for TIR1/AFB-CNGC14-mediated Ca^2+^ signaling in the auxin-regulated vacuolar trafficking of PIN proteins, that depends on TIR1/AFB function (Baster et al., 2013), or PIN polarization (Sauer et al., 2006; Prat et al., 2018; Mazur et al., 2020; Mazur et al., 2020). The contrasting results obtained using mutants versus the currently available Ca^2+^ pharmacology or manipulating Ca^2+^ availability illustrate that Ca^2+^ signaling in plants is a highly interconnected system, not allowing for easy, specific manipulations. Instead, the current lack of highly specific pharmacology imply that the most reliable conclusions can be drawn through genetic analysis.

## CONCLUSION

Auxin is well-known to trigger a Ca^2+^ response. However, its physiological role remains poorly understood. Here, we evaluated auxin-induced Ca^2+^signaling in the context of regulating internalization of plasma membrane cargoes. Pharmacological evidence indicated its requirement. However, genetics demonstrated unequivocally that NAA’s effect on internalization is independent of auxin-induced Ca^2+^ signaling.

## MATERIALS AND METHODS

### Plant Growth Conditions

Arabidopsis thaliana seeds were sterilized by using bleach gas (8mL concentrated HCl to 150mL bleach) overnight, afterwards the seeds were sown on Petri dishes (12 cm X 12 cm) containing sterile half-strength Murashige and Skoog (½ MS) medium (0.5 x MS salts, 0.8% sucrose, 0.5 g/L 2-(N-morpholino) ethanesulfonic acid, pH 5.7, and 1% w/v agar), and grown under continuous light (21°C, continuous light), after 2 days vernalization at 4°C in the dark. To prepare 2L liquid CaMIN medium following components were dissolved in MilliQ: 100mL MS basal salt micronutrient solution (catalog Nr), 20 g sucrose, 0.20 g myoinositol, 1.00 g MES, 1.652g NH_4_NO_3_, 0.180g MgSO_4_, 1.920g KNO_3_, 0.152g KH_2_PO_4_ and pH 5.7. The liquid CaPLUS medium contains an additional 0.332g CaCl_2_ per 2L.

### Chemicals

The following hormones/chemicals were used: 10μM NAA (catalog Nr N0903; Duchefa Biochemie). 25μM Brefeldin A (BFA, catalog Nr B6542-25MG). All hormones/drugs were dissolved in 100% dimethylsulfoxide (DMSO; catalog Nr D4540-500ML) and obtained from Sigma Aldrich.

### Plant Lines Used

The Arabidopsis used as control in this study was Columbia (Col-0) ecotype. We used the following mutants and transgenic lines were described previously: *tir1-1* (Ruegger et al., 1998), *afb2-3* (Parry et al., 2009), *afb3-4* (Parry et al., 2009), *tir1-1afb2-3* (Parry et al., 2009), *tir1-1afb3-4* (Parry et al., 2009), *tir1-1afb1-3afb3-4* (Parry et al., 2009), *tir1-1afb2-3afb3-4* (Parry et al., 2009), *yucQ* (Chen et al., 2014), *cngc14-1* and *cngc14-1* (Shih et al., 2015), *cngc6,9, cngc6,14, cngc9,14* and *cngc6,9,14* (Brost et al., 2019), WAVE131-YFP (Geldner et al., 2009), R-GECO1 (Keinath et al., 2015), NES-YC3.6 (Krebs et al., 2012), PM-YC3.6 (Krebs et al., 2012), CRT-D4ER (Bonza et al., 2013), 4mt-YC3.6 (Loro et al., 2012). The NES-YC3.60 (Krebs et al., 2012) reporter was transformed directly into *tir1-1afb2-3afb3-4* (Parry et al., 2009) and *cngc14-1* (Shih et al., 2015). Transformants were selected based on uniform expression, and expression levels comparable to the control NES-YC3.60. Analyses of the Ca^2+^ response was done on T2 generation seedling roots showing strong and uniform expression.

### Immunodetection and Confocal Microscopy

The seedlings used for Immunodetection are 3-day-old and pre-treated in liquid CaPLUS, CaMIN in presence or absence of chemicals as indicated for 30 minutes. The samples were fixed by paraformaldehyde (4%) in PBS for 1 hour in vacuum. The following steps of the immunostaining were performed by the immuno-robot InsituProII, as described by Sauer et al. (2006). The dilutions of the primary antibodies were: goat anti-PIN1 (1:600) (sc-27163, SantaCruz), rabbit anti-PIN2 (1:600) (Abas et al., 2006) and anti-AUX1 (1:400) (AS 12 1868, Agrisera). The secondary antibodies used were AlexaFluor488 donkey anti-goat (1:600) (A-11055, ThermoFisher) and AlexaFluor555 donkey anti-rabbit (1:600) (A-31572, ThermoFisher).

### Microscopy and image analysis

R-GECO1-based Ca^2+^ imaging experiments and analysis were performed as described (Himschoot et al., 2018). Yellow Cameleon-based experiments were performed and analyzed as described (Behera et al., 2013). Confocal laser scanning microscopes Leica SP2 (Leica) or Zeiss 710 CLSM microscopes were used to analyze the immunolocalisations, and imaging of R-GECO1. Fluorescence emission of Alexa488 (ex 488 nm/em 500-545nm), Alexa555 (ex 561nm/em 555-610nm), YFP (ex 514nm/em 520-565nm), was detected using a 63x water objective. Images were analyzed using Fiji (Schindelin et al., 2015). Fiji was used to rotate, crop images and label the region of all the roots for quantification. The proportion of cells with BFA bodies was scored manually and calculated by using Excel. The BoxPlotR was used to generate the box plots figures (Spitzer et al., 2014).

### Statistical analysis

For statistical analysis of the Ca^2+^ imaging data, unpaired two-tailed t-tests with Welch correction for unequal standard deviations between populations where performed using GraphPad Prism (GraphPad Prism 8 for Windows 64-bit, version 8.4.1).

For statistical analysis of the immunolocalization experiments, a logistic regression was performed to compare the presence of BFA bodies in root cells of treated versus untreated roots or wild type versus mutant. A random effect was added to the model for the experiments with multiple repeats to take into account the correlation between measurements done at the same time. The analysis was performed with the glimmix procedure from SAS (Version 9.4 of the SAS System for windows 7 64bit. Copyright 2002-2012 SAS Institute Inc. Cary, NC, USA (www.sas.com)). Maximum likelihood estimation was done with the default estimation method. A Wald-type test was performed to estimate the treatment/genotype effect on the presence of BFA bodies in the root cells.

## Acknowledgements

We thank Yunde Zhao (UC San Diego, USA), Marc Estelle (UC San Diego, USA), Petra Dietrich (F.A. University of Erlangen-Nurnberg, Germany) and Niko Geldner (University of Lausanne, Switserland) for sharing published materials, and NASC for providing seeds. We thank Veronique Storme for help with the statistical analyses. This work was supported by grants of the China Scholarschip Counsil (CSC) (to R.W. and J.C.), the special research fund Ghent University (to E.H.), the European Research Counsil (ERC) (to D.V.D and J.F.), the Deutsche Forschungsgemeinschaft (DFG) through Grants within FOR964 (M.K. and K.S.), Piano di Sviluppo di Ateneo 2019 (University of Milan) (to A.C.), and by a PhD fellowship from the University of Milan (to MG).

## References

Abas L, Benjamins R, Malenica N, Paciorek T, Wisniewska J, Moulinier-Anzola JC, Sieberer T, Friml J, Luschnig C (2006) Intracellular trafficking and proteolysis of the Arabidopsis auxin-efflux facilitator PIN2 are involved in root gravitropism. Nat Cell Biol 8: 249–256

Baster P, Robert S, Kleine-Vehn J, Vanneste S, Kania U, Grunewald W, De Rybel B, Beeckman T, Friml J (2013) SCF(TIR1/AFB)-auxin signalling regulates PIN vacuolar trafficking and auxin fluxes during root gravitropism. EMBO J 32: 260–274

Behera S, Krebs M, Loro G, Schumacher K, Costa A, Kudla J (2013) Ca2+ imaging in plants using genetically encoded Yellow Cameleon Ca2+ indicators. Cold Spring Harb Protoc 2013: 700–703

Bonza MC, Loro G, Behera S, Wong A, Kudla J, Costa A (2013) Analyses of Ca2+ accumulation and dynamics in the endoplasmic reticulum of Arabidopsis root cells using a genetically encoded Cameleon sensor. Plant Physiol 163: 1230–1241

Brost C, Studtrucker T, Reimann R, Denninger P, Czekalla J, Krebs M, Fabry B, Schumacher K, Grossmann G, Dietrich P (2019) Multiple cyclic nucleotide-gated channels coordinate calcium oscillations and polar growth of root hairs. Plant J 99: 910–923

Cao M, Chen R, Li P, Yu Y, Zheng R, Ge D, Zheng W, Wang X, Gu Y, Gelova Z, Friml J, Zhang H, Liu R, He J, Xu T (2019) TMK1-mediated auxin signalling regulates differential growth of the apical hook. Nature 568: 240–243

Chen Q, Dai X, De-Paoli H, Cheng Y, Takebayashi Y, Kasahara H, Kamiya Y, Zhao Y (2014) Auxin overproduction in shoots cannot rescue auxin deficiencies in Arabidopsis roots. Plant Cell Physiol 55: 1072–1079

Cosgrove DJ, Anderson CT (2020) Plant Cell Growth: Do Pectins Drive Lobe Formation in Arabidopsis Pavement Cells? Curr Biol 30: R660–R662

Costa A, Navazio L, Szabo I (2018) The contribution of organelles to plant intracellular Calcium signalling. J Exp Bot

De Vriese K, Costa A, Beeckman T, Vanneste S (2018) Pharmacological Strategies for Manipulating Plant Ca(2+) Signalling. Int J Mol Sci 19

De Vriese K, Himschoot E, Dunser K, Nguyen L, Drozdzecki A, Costa A, Nowack MK, Kleine-Vehn J, Audenaert D, Beeckman T, Vanneste S (2019) Identification of Novel Inhibitors of Auxin-Induced Ca(2+) Signaling via a Plant-Based Chemical Screen. Plant Physiol 180: 480–496

Demidchik V, Maathuis F, Voitsekhovskaja O (2018) Unravelling the plant signalling machinery: an update on the cellular and genetic basis of plant signal transduction. Funct Plant Biol 45: 1–8

Dharmasiri N, Dharmasiri S, Estelle M (2005) The F-box protein TIR1 is an auxin receptor. Nature 435: 441–445

Dindas J, Scherzer S, Roelfsema MRG, von Meyer K, Muller HM, Al-Rasheid KAS, Palme K, Dietrich P, Becker D, Bennett MJ, Hedrich R (2018) AUX1-mediated root hair auxin influx governs SCF(TIR1/AFB)-type Ca(2+) signaling. Nat Commun 9: 1174

Edel KH, Marchadier E, Brownlee C, Kudla J, Hetherington AM (2017) The Evolution of Calcium-Based Signalling in Plants. Curr Biol 27: R667–R679

Fendrych M, Akhmanova M, Merrin J, Glanc M, Hagihara S, Takahashi K, Uchida N, Torii KU, Friml J (2018) Rapid and reversible root growth inhibition by TIR1 auxin signalling. Nat Plants 4: 453–459

Feng W, Kita D, Peaucelle A, Cartwright HN, Doan V, Duan Q, Liu MC, Maman J, Steinhorst L, Schmitz-Thom I, Yvon R, Kudla J, Wu HM, Cheung AY, Dinneny JR (2018) The FERONIA Receptor Kinase Maintains Cell-Wall Integrity during Salt Stress through Ca(2+) Signaling. Curr Biol 28: 666–675 e665

Gallei M, Luschnig C, Friml J (2020) Auxin signalling in growth: Schrodinger’s cat out of the bag. Curr Opin Plant Biol 53: 43–49

Gao Y, Zhang Y, Zhang D, Dai X, Estelle M, Zhao Y (2015) Auxin binding protein 1 (ABP1) is not required for either auxin signaling or Arabidopsis development. Proc Natl Acad Sci U S A 112: 2275–2280

Geldner N, Denervaud-Tendon V, Hyman DL, Mayer U, Stierhof YD, Chory J (2009) Rapid, combinatorial analysis of membrane compartments in intact plants with a multicolor marker set. Plant J 59: 169–178

Guo J, Yang Z (2020) Exocytosis and endocytosis: coordinating and fine-tuning the polar tip growth domain in pollen tubes. J Exp Bot 71: 2428–2438

Himschoot E, Krebs M, Costa A, Beeckman T, Vanneste S (2018) Calcium Ion Dynamics in Roots: Imaging and Analysis. Methods Mol Biol 1761: 115–130

Himschoot E, Pleskot R, Van Damme D, Vanneste S (2017) The ins and outs of Ca(2+) in plant endomembrane trafficking. Curr Opin Plant Biol 40: 131–137

Huang R, Zheng R, He J, Zhou Z, Wang J, Xiong Y, Xu T (2019) Noncanonical auxin signaling regulates cell division pattern during lateral root development. Proc Natl Acad Sci U S A 116: 21285–21290

Keinath NF, Waadt R, Brugman R, Schroeder JI, Grossmann G, Schumacher K, Krebs M (2015) Live Cell Imaging with R-GECO1 Sheds Light on flg22- and Chitin-Induced Transient [Ca(2+)]cyt Patterns in Arabidopsis. Mol Plant 8: 1188–1200

Kepinski S, Leyser O (2005) The Arabidopsis F-box protein TIR1 is an auxin receptor. Nature 435: 446–451

Konrad KR, Maierhofer T, Hedrich R (2018) Spatio-temporal Aspects of Ca2+ Signalling: Lessons from Guard Cells and Pollen Tubes. J Exp Bot

Krebs M, Held K, Binder A, Hashimoto K, Den Herder G, Parniske M, Kudla J, Schumacher K (2012) FRET-based genetically encoded sensors allow high-resolution live cell imaging of Ca(2)(+) dynamics. Plant J 69: 181–192

Kudla J, Becker D, Grill E, Hedrich R, Hippler M, Kummer U, Parniske M, Romeis T, Schumacher K (2018) Advances and current challenges in calcium signaling. New Phytol 218: 414–431

Lavy M, Estelle M (2016) Mechanisms of auxin signaling. Development 143: 3226–3229

Li R, Sun R, Hicks GR, Raikhel NV (2015) Arabidopsis ribosomal proteins control vacuole trafficking and developmental programs through the regulation of lipid metabolism. Proc Natl Acad Sci U S A 112: E89–98

Li T, Yan A, Bhatia N, Altinok A, Afik E, Durand-Smet P, Tarr PT, Schroeder JI, Heisler MG, Meyerowitz EM (2019) Calcium signals are necessary to establish auxin transporter polarity in a plant stem cell niche. Nat Commun 10: 726

Loro G, Drago I, Pozzan T, Schiavo FL, Zottini M, Costa A (2012) Targeting of Cameleons to various subcellular compartments reveals a strict cytoplasmic/mitochondrial Ca(2)(+) handling relationship in plant cells. Plant J 71: 1–13

Mazur E, Gallei M, Adamowski M, Han H, Robert HS, Friml J (2020) Clathrin-mediated trafficking and PIN trafficking are required for auxin canalization and vascular tissue formation in Arabidopsis. Plant Sci 293: 110414

Mazur E, Kulik I, Hajny J, Friml J (2020) Auxin canalization and vascular tissue formation by TIR1/AFB-mediated auxin signaling in Arabidopsis. New Phytol 226: 1375–1383

Nagai T, Yamada S, Tominaga T, Ichikawa M, Miyawaki A (2004) Expanded dynamic range of fluorescent indicators for Ca(2+) by circularly permuted yellow fluorescent proteins. Proc Natl Acad Sci U S A 101: 10554–10559

Nemhauser JL (2018) Back to basics: what is the function of an Aux/IAA in auxin response? New Phytol 218: 1295–1297

Paciorek T, Zazimalova E, Ruthardt N, Petrasek J, Stierhof YD, Kleine-Vehn J, Morris DA, Emans N, Jurgens G, Geldner N, Friml J (2005) Auxin inhibits endocytosis and promotes its own efflux from cells. Nature 435: 1251–1256

Pan J, Fujioka S, Peng J, Chen J, Li G, Chen R (2009) The E3 ubiquitin ligase SCFTIR1/AFB and membrane sterols play key roles in auxin regulation of endocytosis, recycling, and plasma membrane accumulation of the auxin efflux transporter PIN2 in Arabidopsis thaliana. The Plant cell 21: 568–580

Parry G, Calderon-Villalobos LI, Prigge M, Peret B, Dharmasiri S, Itoh H, Lechner E, Gray WM, Bennett M, Estelle M (2009) Complex regulation of the TIR1/AFB family of auxin receptors. Proc Natl Acad Sci U S A 106: 22540–22545

Pierre-Jerome E, Moss BL, Lanctot A, Hageman A, Nemhauser JL (2016) Functional analysis of molecular interactions in synthetic auxin response circuits. Proc Natl Acad Sci U S A 113: 11354–11359

Platre MP, Bayle V, Armengot L, Bareille J, Marques-Bueno MDM, Creff A, Maneta-Peyret L, Fiche JB, Nollmann M, Miege C, Moreau P, Martiniere A, Jaillais Y (2019) Developmental control of plant Rho GTPase nano-organization by the lipid phosphatidylserine. Science 364: 57–62

Powers SK, Strader LC (2020) Regulation of auxin transcriptional responses. Dev Dyn 249: 483–495

Prat T, Hajny J, Grunewald W, Vasileva M, Molnar G, Tejos R, Schmid M, Sauer M, Friml J (2018) WRKY23 is a component of the transcriptional network mediating auxin feedback on PIN polarity. PLoS Genet 14: e1007177

Robert S, Kleine-Vehn J, Barbez E, Sauer M, Paciorek T, Baster P, Vanneste S, Zhang J, Simon S, Covanova M, Hayashi K, Dhonukshe P, Yang Z, Bednarek SY, Jones AM, Luschnig C, Aniento F, Zazimalova E, Friml J (2010) ABP1 mediates auxin inhibition of clathrin-dependent endocytosis in Arabidopsis. Cell 143: 111–121

Roosjen M, Paque S, Weijers D (2018) Auxin Response Factors: output control in auxin biology. J Exp Bot 69: 179–188

Ruegger M, Dewey E, Gray WM, Hobbie L, Turner J, Estelle M (1998) The TIR1 protein of Arabidopsis functions in auxin response and is related to human SKP2 and yeast grr1p. Genes Dev 12: 198–207

Sauer M, Balla J, Luschnig C, Wisniewska J, Reinohl V, Friml J, Benkova E (2006) Canalization of auxin flow by Aux/IAA-ARF-dependent feedback regulation of PIN polarity. Genes Dev 20: 2902–2911

Schindelin J, Rueden CT, Hiner MC, Eliceiri KW (2015) The ImageJ ecosystem: An open platform for biomedical image analysis. Mol Reprod Dev 82: 518–529

Shih HW, DePew CL, Miller ND, Monshausen GB (2015) The Cyclic Nucleotide-Gated Channel CNGC14 Regulates Root Gravitropism in Arabidopsis thaliana. Curr Biol 25: 3119–3125

Spitzer M, Wildenhain J, Rappsilber J, Tyers M (2014) BoxPlotR: a web tool for generation of box plots. Nat Methods 11: 121–122

Stael S, Wurzinger B, Mair A, Mehlmer N, Vothknecht UC, Teige M (2012) Plant organellar calcium signalling: an emerging field. J Exp Bot 63: 1525–1542

Tian W, Hou C, Ren Z, Wang C, Zhao F, Dahlbeck D, Hu S, Zhang L, Niu Q, Li L, Staskawicz BJ, Luan S (2019) A calmodulin-gated calcium channel links pathogen patterns to plant immunity. Nature 572: 131–135

Vanneste S, Friml J (2013) Calcium: The Missing Link in Auxin Action. Plants (Basel) 2: 650–675

Waadt R, Krebs M, Kudla J, Schumacher K (2017) Multiparameter imaging of calcium and abscisic acid and high-resolution quantitative calcium measurements using R-GECO1-mTurquoise in Arabidopsis. New Phytol 216: 303–320

Wang Q, Qin G, Cao M, Chen R, He Y, Yang L, Zeng Z, Yu Y, Gu Y, Xing W, Tao WA, Xu T (2020) A phosphorylation-based switch controls TAA1-mediated auxin biosynthesis in plants. Nat Commun 11: 679

Xu T, Wen M, Nagawa S, Fu Y, Chen JG, Wu MJ, Perrot-Rechenmann C, Friml J, Jones AM, Yang Z (2010) Cell surface- and rho GTPase-based auxin signaling controls cellular interdigitation in Arabidopsis. Cell 143: 99–110

Zhang J, Mazur E, Balla J, Gallei M, Kalousek P, Medvedova Z, Li Y, Wang Y, Prat T, Vasileva M, Reinohl V, Prochazka S, Halouzka R, Tarkowski P, Luschnig C, Brewer PB, Friml J (2020) Strigolactones inhibit auxin feedback on PIN-dependent auxin transport canalization. Nat Commun 11: 3508

Zhang J, Vanneste S, Brewer PB, Michniewicz M, Grones P, Kleine-Vehn J, Lofke C, Teichmann T, Bielach A, Cannoot B, Hoyerova K, Chen X, Xue HW, Benkova E, Zazimalova E, Friml J (2011) Inositol trisphosphate-induced Ca2+ signaling modulates auxin transport and PIN polarity. Dev Cell 20: 855–866

Zhao Y, Araki S, Wu J, Teramoto T, Chang YF, Nakano M, Abdelfattah AS, Fujiwara M, Ishihara T, Nagai T, Campbell RE (2011) An expanded palette of genetically encoded Ca(2)(+) indicators. Science 333: 1888–1891

